# GenVarLoader: An accelerated dataloader for applying deep learning to personalized genomics

**DOI:** 10.1101/2025.01.15.633240

**Authors:** David Laub, Aaron Ho, Jeff Jaureguy, Adam Klie, Rany M. Salem, Graham McVicker, Hannah Carter

**Affiliations:** Bioinformatics and Systems Biology Program, University of California, San Diego, La Jolla, CA, 92093; Integrative Biology Laboratory, Salk Institute for Biological Studies, La Jolla, CA, 92093; Herbert Wertheim School of Public Health and Longevity Science, University of California, San Diego, La Jolla, CA, 92093; Department of Medicine, Division of Genomics and Precision Medicine, University of California, San Diego, La Jolla, CA, 92093; Moores Cancer Center, University of California, San Diego, La Jolla, CA, 92093

## Abstract

Deep learning sequence models trained on personalized genomics can improve variant effect prediction, however, applications of these models are limited by computational requirements for storing and reading large datasets. We address this with GenVarLoader, which stores personalized genomic data in new memory-mapped formats with optimal data locality to achieve ∼1,000x faster throughput and ∼2,000x better compression compared to existing alternatives.

## Main

Personalized sequence models can inform precision medicine, leading to more accurate gene regulatory predictions and improved understanding of variant driven disease risk^1^. Deep learning (DL) sequence models trained on reference genome sequences to predict functional data have improved our ability to detect the effects of noncoding variants on gene expression^2^, transcription factor binding^3^, histone modifications^2^, chromatin accessibility^4^, and DNA contact maps^5^. However, these models fail to explain individual variation in gene expression^6,7^. Recent studies address this by training sequence models on personalized genomes paired with gene expression^8,9^. As they improve, DL sequence models may complement current methods for imputing functional data for phenotypic association studies (e.g. transcriptome-wide association studies) and for predicting phenotypic data directly (e.g. polygenic risk scores). This sets the stage for the application of sequence models to personalized data from biobanks such as UK BioBank^10^ and All Of Us (AoU)^11^, which contain hundreds of thousands of individuals’ genomic and phenotypic data. However application of DL sequence models to personalized genomes is currently extremely computationally-intensive due to the petabyte-scale of personalized genomic data and slow read throughput.

Efficient methods for generating personalized genomes and functional tracks are essential to scale the training and application of DL sequence models. Most approaches to train on personalized genomes have used bcftools^12^ to reconstruct diploid genomes as FASTA files after taking a reference genome FASTA and variant call format (VCF) files as input^8,9^. Similarly, RNA-seq read depth is typically stored in BigWig format with regions specified in BED files. However, this process demands excessive storage, limits throughput, and lacks support for indels. Writing personal genomes as FASTA files inflates storage requirements by orders of magnitude compared to VCF files. For example, storing the one million genomes from the AoU project would require 1.97 petabytes of storage and cost over $50,000 per month (*Methods*). Additionally, FASTA-based data loading limits throughput, lowers GPU utilization, inflates compute costs, and delays model iteration. Finally, supporting indels also requires re-aligning functional tracks and no published tools currently support this.

Here, we introduce GenVarLoader (GVL), which generates personalized genomes and functional tracks on-the-fly, supports indels, and achieves throughput up to 1,000 times faster than alternatives. By eliminating data loading bottlenecks, GVL ensures efficient model training and inference. Additionally, GVL is scalable, with sustained performance on datasets of hundreds of thousands of genomes.

To easily integrate with common workflows, GVL takes VCF, BigWig, and BED files as input and yields a PyTorch compatible dataloader (**Fig. 1A**). GVL supports indels by re-aligning the read depth to changes in genome length and adjusting the coordinates for regions of interest. In addition, it reconstructs personalized genomes and re-aligned read depth on-the-fly (**Fig. 1A**).

**Figure 1.**
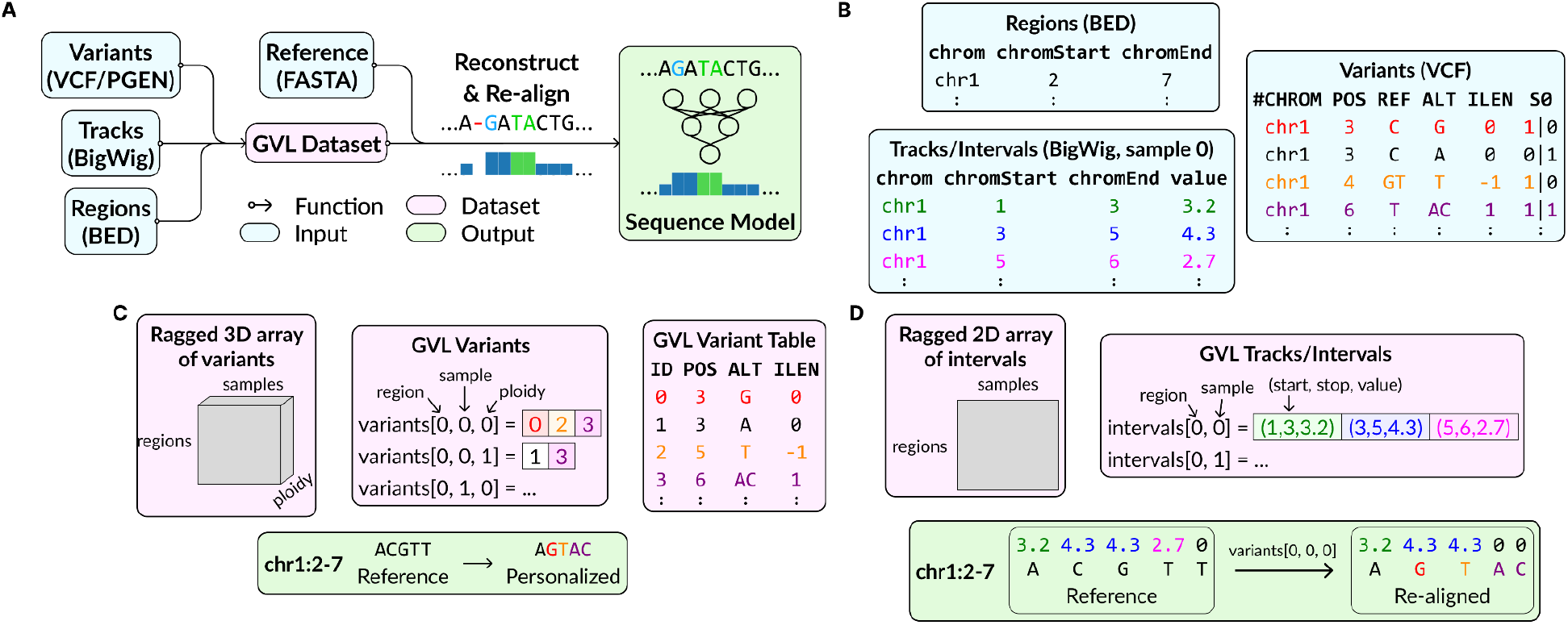
**A)** GenVarLoader (GVL) workflow. Variants, read depth, and regions of interest are converted into a GVL dataset. The GVL dataset and a reference genome are used during training and inference to reconstruct personalized sequences and re-align tracks for sequence models. Tracks are re-aligned to support indels, skipping values overlapping deletions and duplicating values for insertions. **B)** Example of source data and how **C)** variants and **D)** tracks/intervals are represented in a GVL dataset and used in personalized and re-aligned sequences and tracks.

To optimize throughput, GVL reorganizes variants and BigWig files into memory-mapped sample-major layouts. In contrast, VCF and PGEN files store genotypes as dense, variant-major matrices, which are inefficient for randomly sampling regions across individuals (**Fig. 1B**). These formats are also block-compressed and indexed, requiring decompression and search for random access. Similarly, BigWig files store data per sample, needing expensive operating system calls when a different file is accessed (**Fig. 1B**). Instead, GVL sparsifies genotypes matrices and rearranges data as memory-mapped ragged arrays (**Fig. 1C-D**). This eliminates the need for decompression and search, optimizes data locality, and reduces I/O overhead resulting in substantially faster data retrieval.

We benchmarked GVL’s storage footprint and sequence throughput using three datasets of varying size: (1) 62 whole genome sequencing (WGS) and ATAC-seq samples from The Cancer Genome Atlas (TCGA)^13,14^; (2) 3,202 WGS samples from the 1,000 Genomes Project; (3) 487,409 imputed genotype samples from the UK BioBank limited to chromosome 22. We compared GVL’s storage footprint to storing personalized genomes as two compressed FASTA files, corresponding to prior approaches^8,9^. For the 1,000 Genomes Project^15^, personalized FASTA files required 6.3 TB whereas GVL required only 3.1 GB, an over 2,000-fold reduction, with similar results for the UK BioBank^10^ (**Fig. 2A**). We compared the maximum throughput of GVL to reading FASTA files from bcftools using up to 64 CPUs and 16 GB of total RAM. To reflect common lengths used in practice and to assess different data access patterns, we tested sequence lengths ranging from 2 kbp to 1 Mbp. GVL achieved a 300-1,000x speed-up over reading FASTA and exceeded the maximum input bandwidth of modern GPUs such as the NVIDIA A100 (**Fig. 2B**). We found that GVL had consistent throughput for all three datasets, demonstrating its scalability to hundreds of thousands of samples or more.

**Figure 2.**
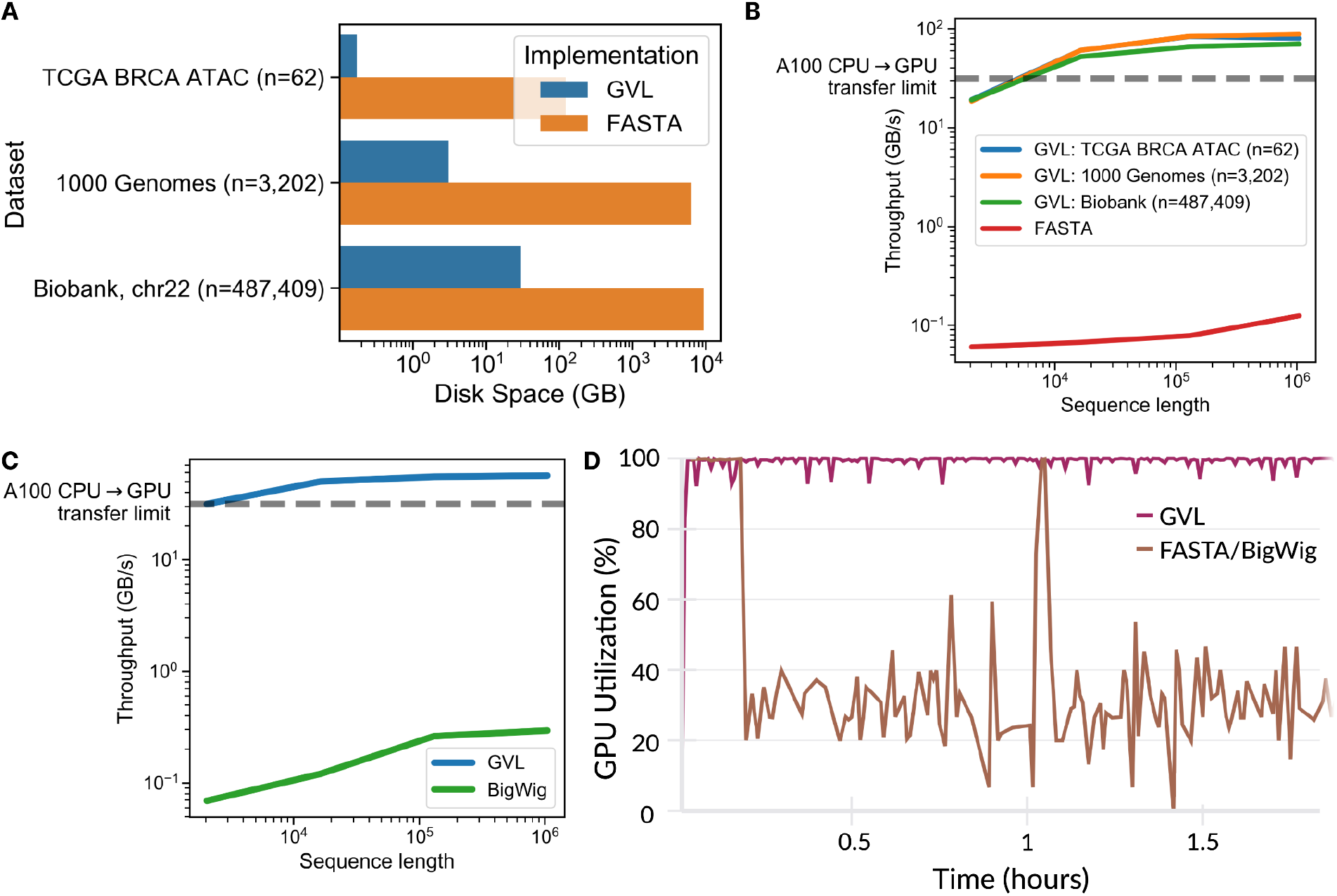
Advantages of GenVarLoader (GVL) over alternatives concerning storage size and data loading throughput. **A)** Storage requirements for personalized genomes in GVL compared to compressed FASTA files. **B)** GVL throughput for reconstructing haplotypes versus reading FASTA from bcftools; GVL is 300-1,000 times faster. **C)** Throughput for generating re-aligned tracks with GVL versus reading BigWig files; GVL is 190-450 times faster. **D)** GPU utilization during training using GVL versus standard tools; GVL enabled full GPU utilization, a 3x improvement.

Similarly, we benchmarked GVL’s performance for generating functional track data re-aligned to personal genomes. Using the same resources as before (up to 64 CPUs and 16 GB total RAM), we compared GVL against a parallelized implementation of pyBigWig on the TCGA dataset. Testing sequence lengths ranging from 2 kbp to 1 Mbp, we observed 190-450x faster throughput compared to the pyBigWig implementation (**Fig. 2C)**. Once again, GVL’s throughput exceeded modern GPU input bandwidth.

We next tested whether GVL could eliminate data loading bottlenecks during model training by achieving full GPU utilization. Using the TCGA ATAC-seq dataset, 64 CPUs, 8 GB total RAM, and one A30 GPU, we compared GVL to a data loading pipeline built with pysam^16^ and pyBigWig^17^. GVL sustained full GPU utilization while pysam and pyBigWig reduced GPU utilization to ∼ 30% (**Fig. 2D**).

GenVarLoader addresses critical barriers in scaling sequence models to personalized genomics. By reducing storage requirements, adding indel support, and improving throughput by up to 1,000-fold over existing tools, GVL enables efficient training and inference with DL sequence models on biobank-scale genomic datasets. Future work will support alternative variant formats (e.g. Hail^18^, TileDB-VCF^19^) and variant calls from pangenome and multiple references^20^. Finally, as GPU hardware evolves, further optimizations can be pursued to ensure GPU bandwidth saturation.

## Methods

### Benchmarking datasets

We benchmarked GVL utilizing three datasets encoded as VCF, PGEN, or BigWig files: (1) paired WGS and ATAC-seq data for 62 breast cancer samples from TCGA representing 4,492,345 unique variants^13,14^ ;(2) WGS data for 3,202 individuals from the 1,000 Genomes Project^15^ containing 73,452,337 unique variants; (3) imputed genotypes for 478,409 samples were obtained from the UK BioBank consisting of 1,247,115 unique variants restricted to chromosome 22^10^.

### Benchmarking approach

To compare the storage requirements of GenVarLoader (GVL) to a bcftools consensus-based approach (denoted FASTA), we estimated the size needed by bcftools as the compressed GRCh37 reference genome (987 MB) multiplied by twice the number of individuals for diploid genomes. We measured GVL’s storage using dust^21^ v1.1.1.

To measure throughput, we ran benchmarks on a SLURM-based high-performance computing cluster allocating 16 GB of total RAM and 1 to 64 virtual CPUs from an AMD EPYC 7543 processor. We selected genome-wide non-overlapping intervals at sequence lengths of 2,048, 16,384, 131,072, and 1,048,576 nucleotides. Batch sizes were chosen at powers of two ensuring that batches did not exceed 8 GB. Each benchmarking run proceeded until 2^29^ base pairs were processed or 10 batches were completed, whichever was larger, and each configuration was repeated 5 times. The maximum throughput measured across threads, batch sizes, and replicates is shown in figures 1B and C. For reading FASTA and BigWig files, we used pysam v0.22.1 and pyBigWig v0.3.23, respectively.

To assess GPU utilization, we trained a convolutional neural network on one-hot encoded DNA sequences of 2,048 bp to predict 1,048 bp of ATAC-seq read depth centered at peaks. The model architecture is adapted from BPNet^3^ using an initial convolutional layer with a kernel size of 21, followed by 8 dilated convolutional layers. Predictions for each haplotype were summed together to make the final prediction, similar to existing approaches^9,22^. Each training run was allocated 32 CPUs, 8 GB RAM, and one NVIDIA A30 GPU. GPU utilization was measured with Weights & Biases^23^.

### Estimating cloud storage costs

We estimated cloud storage costs for 1 million genomes stored as compressed FASTA files using the AoU pricing of $0.026 per GB-month and the GRCh37 reference genome size (987 MB). Each genome requires one FASTA file for each haplotype, totaling 1,000,000 genomes x 0.987 GB/file x 2 files/genome x $0.026 per GB-month = $51,324 per month.

## Data availability

WGS VCFs and ATAC-seq BigWigs for TCGA breast cancer samples are available at the Genomic Data Commons portal https://portal.gdc.cancer.gov/. WGS variant data from the 1,000 Genomes Project are available at https://www.internationalgenome.org/data-portal/data-collection/30x-grch38. Imputed UK Biobank data were obtained under project ID 37671 and data are available to qualified investigators at https://www.ukbiobank.ac.uk/. GenVarLoader datasets used for benchmarking on the 1,000 Genomes Project are available on Zenodo^24^.

## Code availability

GVL is available under the MIT license as a python package https://pypi.org/project/genvarloader/ with documentation at https://genvarloader.readthedocs.io/en/latest/. The results presented here correspond to GVL v0.6.1 and, along with benchmarking and plotting code, is archived at Zenodo^25^. The latter is also available as a GitHub repository at https://github.com/d-laub/gvl-paper.

## Acknowledgements

This work was supported by NIH-NHGRI grant HG011315 to GM, the Frederick B. Rentschler Developmental Chair to GM, NIH-NHGRI F31 fellowship HG013262 to JJ, NIH grant 2P41GM103504 for infrastructure, and NIH-NHGRI 1U01HG012059 to HC. We would like to thank Brad Balderson for his feedback on the manuscript.

## Author contributions

DL designed and implemented GenVarLoader (GVL), and wrote benchmarking code and the manuscript. AH implemented the first iteration of GVL. JJ tested several iterations of GVL. AK helped implement the neural network used for benchmarking GPU utilization. RS provided UK BioBank data. GM and HC supervised the work. All authors read and corrected the final manuscript.

## Ethics declarations

The authors declare no competing interests.

## References

1. He, A. Y., Palamuttam, N. P. & Danko, C. G. Training deep learning models on personalized genomic sequences improves variant effect prediction. 2024.10.15.618510 Preprint at 10.1101/2024.10.15.618510 (2024).

2. Avsec, Ž. et al. Effective gene expression prediction from sequence by integrating long-range interactions. Nat. Methods 18, 1196–1203 (2021).

3. Avsec, Ž. et al. Base-resolution models of transcription-factor binding reveal soft motif syntax. Nat. Genet. 53, 354–366 (2021).

4. Trevino, A. E. et al. Chromatin and gene-regulatory dynamics of the developing human cerebral cortex at single-cell resolution. Cell 184, 5053-5069.e23 (2021).

5. Zhou, J. Sequence-based modeling of three-dimensional genome architecture from kilobase to chromosome scale. Nat. Genet. 54, 725–734 (2022).

6. Huang, C. et al. Personal transcriptome variation is poorly explained by current genomic deep learning models. Nat. Genet. 55, 2056–2059 (2023).

7. Sasse, A. et al. Benchmarking of deep neural networks for predicting personal gene expression from DNA sequence highlights shortcomings. Nat. Genet. 55, 2060–2064 (2023).

8. Drusinsky, S., Whalen, S. & Pollard, K. S. Deep-learning prediction of gene expression from personal genomes. 2024.07.27.605449 Preprint at 10.1101/2024.07.27.605449 (2024).

9. Rastogi, R., Reddy, A. J., Chung, R. & Ioannidis, N. M. Fine-tuning sequence-to-expression models on personal genome and transcriptome data. 2024.09.23.614632 Preprint at 10.1101/2024.09.23.614632 (2024).

10. Sudlow, C. et al. UK Biobank: An Open Access Resource for Identifying the Causes of a Wide Range of Complex Diseases of Middle and Old Age. PLOS Med. 12, e1001779 (2015).

11. The All of Us Research Program Investigators. The “All of Us” Research Program. N. Engl. J. Med. 381, 668–676 (2019).

12. Danecek, P. et al. Twelve years of SAMtools and BCFtools. GigaScience 10, giab008 (2021).

13. Weinstein, J. N. et al. The Cancer Genome Atlas Pan-Cancer analysis project. Nat. Genet. 45, 1113–1120 (2013).

14. Corces, M. R. et al. The chromatin accessibility landscape of primary human cancers. Science 362, eaav1898 (2018).

15. Auton, A. et al. A global reference for human genetic variation. Nature 526, 68–74 (2015).

16. pysam-developers/pysam. pysam-developers (2024).

17. deeptools/pyBigWig. The deepTools ecosystem (2024).

18. Hail Team, H. T. Hail. (2024).

19. TileDB-Inc/TileDB-VCF. TileDB, Inc. (2024).

20. Chen, N.-C., Solomon, B., Mun, T., Iyer, S. & Langmead, B. Reference flow: reducing reference bias using multiple population genomes. Genome Biol. 22, 8 (2021).

21. andy.boot. bootandy/dust. (2024).

22. Celaj, A. et al. An RNA foundation model enables discovery of disease mechanisms and candidate therapeutics. 2023.09.20.558508 Preprint at 10.1101/2023.09.20.558508 (2023).

23. Biewald, L. Experiment Tracking with Weights and Biases. (2020).

24. Laub, D. GenVarLoader Paper Datasets. Zenodo 10.5281/zenodo.14367502 (2024).

25. Laub, D. GenVarLoader Paper Code. Zenodo 10.5281/zenodo.14375325 (2024).

